# Machine Learning Algorithms for Predicting Coronary Artery Disease: Efforts Toward an Open Source Solution

**DOI:** 10.1101/2020.02.13.948414

**Authors:** Aravind Akella, Vibhor Kaushik

## Abstract

The development of Coronary Artery Disease (CAD), one of the most prevalent diseases in the world, is heavily influenced by several modifiable risk factors. Predictive models built using machine learning (ML) algorithms may assist healthcare practitioners in timely detection of CAD, and ultimately, may improve outcomes. In this study, we have applied six different ML algorithms to predict the presence of CAD amongst patients listed in an openly available dataset provided by the University of California Irvine (UCI) Machine Learning Repository, named “the Cleveland dataset.” All six ML algorithms achieved accuracies greater than 80%, with the “Neural Network” algorithm achieving accuracy greater than 93%. The recall achieved with the “Neural Network” model is also highest of the six models (0.93). Additionally, five of the six algorithms resulted in very similar AUC-ROC curves. The AUC-ROC curve corresponding to the “Neural Network” algorithm is slightly steeper implying higher “true positive percentage” achieved with this model. We also extracted the variables of importance in the “Neural Network” model to help in the risk assessment. We have released the full computer code generated in this study in the public domain as a preliminary effort toward developing an open solution for predicting the presence of coronary artery disease in a given population and present a workflow model for implementing a possible solution.

## Introduction

Coronary artery disease (CAD) is the most common type of heart disease affecting millions worldwide. According to recent statistics from the American Heart Association, CAD accounted for 13% of deaths in the United States in 2016 (see https://healthmetrics.heart.org/wp-content/uploads/2019/02/At-A-Glance-Heart-Disease-and-Stroke-Statistics-%E2%80%93-2019.pdf). Worldwide, CAD was found to be the cause of 15.6% of all deaths in 2015, making it one of the most common causes of mortality[1]. Because CAD is associated with several modifiable risk factors pertaining to lifestyle and intervention, timing of detection and diagnostic accuracy are especially relevant in its treatment. Over the past several years, approaches that include machine learning (ML) are making significant impact in the detection and diagnosis of diseases[2, 3, also see 4–7]. In general, the ML approach involves “training” an algorithm with a control dataset for which the “outcome” (disease or no disease) is known, and then applying this trained algorithm to predict outcomes in patients whose CAD status is not yet determined. When applied to disease outcomes involving modifiable risk factors (as in CAD), the ML approach may hasten the diagnostic process while decreasing error, allowing for improved mortality and morbidity.

The application of ML concepts to CAD has been significantly hampered by the availability of appropriate clinical datasets. However, one of the components of the “UCI Heart Disease Dataset”, dubbed as the “Cleveland Dataset”, is publicly available on the UCI Machine Learning Repository (see https://archive.ics.uci.edu/ml/datasets/Heart+Disease). Originally intended to be a teaching aid, the Cleveland dataset has been highly exploited for exploring ML concepts. Available since 1988, the Cleveland dataset has so far received more than one million downloads and is currently ranked as the fifth most popular dataset in the UCI repository. A GoogleScholar search with the terms “Cleveland dataset,” “heart disease,” and “machine learning,” returns little more than 300 records available since 2010, of which most studies analyzed one ML method at a time (for a latest survey review, see[8]). In addition, there is no way to verify claims in any of the publications for the accuracy of the algorithms, as the computer code has not been made publicly available^1^. A PubMed search using the same keywords identified ten original articles and one comprehensive review article[9]. Of the ten original articles, three used the Cleveland dataset, whereas the remaining seven used proprietary datasets. Of the ten, none of the studies have made the computer code publicly available. In our view, all recent studies pertaining to the application of the ML approach to CAD thus far appear exploratory rather than seeking to provide clinical assistance to healthcare practitioners in the treatment of CAD.

We have now undertaken a comparative analysis by applying six different ML algorithms (models) using the UCI Cleveland dataset to predict disease outcomes. In an effort to initiate an open source ML solution for detecting CAD, we have deposited our computer code on the GitHub platform (see https://github.com/aa54/CAD_1), making it available for other researchers to test and improve our work. We also welcome the opportunity to gain access to larger datasets to further our efforts towards an open source solution. As for the six ML algorithms used in our study, we have found that all six (Linear Regression, Regression Tree, Random Forest, Support-Vector Machine, Random Forest, Nearest Neighbor, and k-Nearest Neighbor) perform well with an accuracy of greater than ∼80%, with the Nearest Neighbor’s accuracy greater than 93%. We consider Accuracy, Recall, F1 score, and the Area Under the Curve–Receiver Operating (AUC-ROC) as performance metrics for comparative analysis amongst the ML models.

Numerous risk factor variables contribute to the development of CAD, some of which can be controlled (or modifiable) (e.g., see[10]). These include high blood pressure, high cholesterol, smoking, diabetes, obesity, lack of physical activity, unhealthy diet, and stress. The risk factors that cannot be controlled (or non-modifiable) are age, sex (gender), family history, and race or ethnicity. Traditional approaches assess these risk factors to predict future risk (prognosis) of cardiovascular disease[11]. However, a large number of individuals at risk of cardiovascular disease fail to be identified by these approaches, while some individuals not at risk are given preventive treatment unnecessarily (see [12, 13]). Several of the ML algorithms have the ability to summarize the impact of individual variables on response variable and are referred as “variables of importance”, thus leading to the building of prognostic models[14]. In the present study, we extract the variables of importance in one model in an effort to demonstrate the feasibility of including risk assessment component in the ML based open solution.

## Dataset and Preprocessing

The dataset used in this study is downloaded from the repository maintained by the UCI (University of California, Irvine, CA) Center for Machine Learning and Intelligent Systems. The repository contains four datasets from four different hospitals. The Cleveland dataset contains less missing attributes than the other datasets and has more records. This dataset has fourteen sets of variables on 303 patients. Table 1 lists all fourteen variables in the dataset, the associated datatype for each of the attributes, and a brief description of the attributes.

**Table 1:** Dataset variables names, types, and descriptions

| Variable | Type | Description |
| --- | --- | --- |
| age | continuous | patient age in years |
| sex | categorical | patient gender (1 = male; 0 = female) |
| cp | categorical | chest pain (1 = typical angina; 2 = atypical angina; 3 = non-anginal pain; 4 = no pain) |
| trestbps | continuous | resting blood pressure (in mmHg) on admission to the hospital |
| chol | continuous | serum cholesterol in mg/dL |
| fbs | categorical | fasting blood sugar higher than 120 mg/dL (1 = true; 0 = false) |
| restecg | categorical | resting EKG (0 = normal; 1 = ST-T wave abnormality; 2 = probable/definite LVH) |
| thalach | continuous | maximum heart rate achieved (during thallium test) |
| exang | categorical | exercise induced angina (1 = yes; 0 = no) |
| oldpeak | continuous | ST depression induced by exercise relative to rest |
| slope | categorical | slope of the peak exercise ST segment (1 = up-sloping; 2 = flat; 3 = down-sloping) |
| ca | categorical | number of major vessels (0 to 3) colored by fluoroscopy |
| thal | categorical | thallium heart scan (3 = normal; 6 = fixed defect; 7 = reversible defect) |
| num | categorical | diagnosis of heart disease (angiographic disease status) (0 = absent; 1 to 4 = present) |

During the data preprocessing steps, (1) six variables with unknown values were dropped resulting in a dataset of 297 observations; (2) two dataset columns of factor datatype were converted to numeric datatype; (3) continuous variables (four, excluding age) were normalized; (4) the column “num” was renamed to “hd” for clarity; and (5) the values 1 to 4 in the heart disease (hd) column are replaced with value 1 to form a binary classification—disease (value of 1) or no disease (value of 0).

The correlation matrix for the fourteen variables in the dataset is shown in Figure 1, depicting the Pearson coefficients, or r, corresponding to the association between two variables, which can range from –1 to +1. A higher absolute value of r indicates a stronger correlation between two variables. A positive r indicates a direct relationship between two variables, whereas a negative r suggests an indirect relationship. In this analysis, we considered an absolute value of 0.5 as threshold, that is, if r is greater than 0.5 or less than –0.5, we assume those two variables are correlated. The above correlation matrix shows that only a few variables correlate with coefficients greater than 0.5, demonstrating poor correlation between the variables. Having a low value of r indicates that all these independent variables and can be included in the machine learning model building.

**Figure 1:**
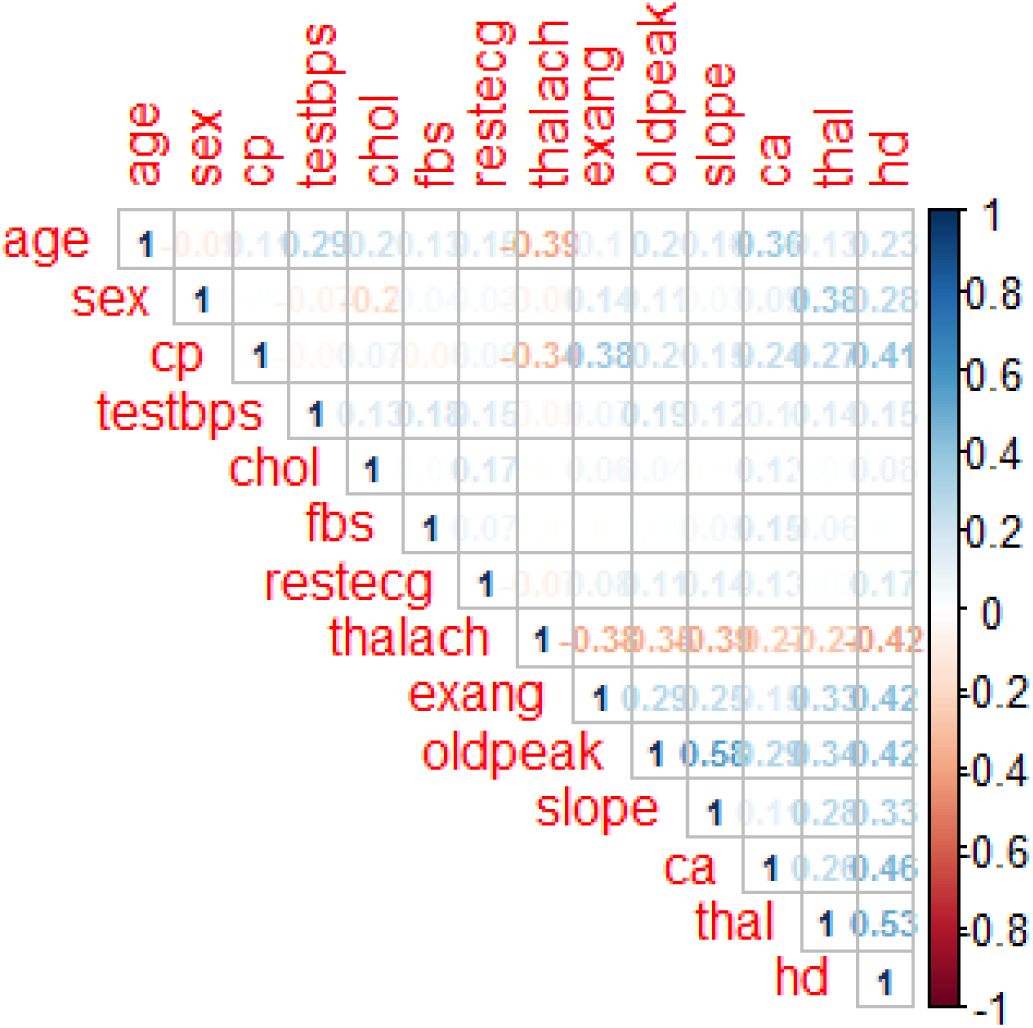
Correlation matrix of the various parameters in the dataset.

Additional analysis of the dataset is presented in the Supplementary Material.

## Machine Learning Models, Model Building, and Model Evaluation

In applying machine learning models, it is generally understood that no single algorithm is superior to the others[15]. In machine learning, if every instance in the dataset is given to the model with known labels (the corresponding correct outputs), like in the Cleveland dataset, then the learning is called “supervised”, in contrast to “unsupervised” learning in which instances are unlabeled. Below we present the general idea on how each of the six supervised machine learning algorithms work on the dataset and any assumptions we make in each case. Each algorithm is first trained (or fitted) with a fraction of the dataset, usually known as the “training set” and then tested on the “test set” that is put aside as “unseen data” for evaluating the algorithm. For a detailed description of the models we refer the reader to the excellent treatise by James et al[16].

### Binary Logistic Regression

In Binary Logistic Regression, each independent variable in every instance of the dataset is multiplied by a weight and the cumulative result is passed to a sigmoid function. The function maps real values into probability values between 0 and 1. In our present modeling, we left the threshold to the default value, which is 0.5, such that for probability values greater than 0.5, the model predicts the dependent variable to be 1 (the patient has CAD), and for values less than or equal to 0.5, the model predicts the dependent variable to be 0 (the patient does not have CAD).

### Decision Tree

Decision (Regression) Tree is tree-like structure that classifies instances by sorting them based on the values of the variables. Each node in a decision tree represents a variable in an instance to be classified, and each branch represents a value that the node can assume. Instances are classified starting at the root node and sorted based on the values of the variables. The variable that best divides the dataset would be the root node of the tree. Internal nodes (or split nodes) are the decision-making part that make a decision, based on multiple algorithms, and to visit subsequent nodes. The split process is terminated when a user defined criteria is reached at the leaf (for the present modeling, we left it to be the default value, which is 20). The paths from root nodes to the leaf nodes represent classification rules.

### Random Forest

Random Forest is ensemble model constating of multiple regression trees like in a forest. Random forest combines several regression trees, trains each one on a slightly different set of the dataset instances, splitting nodes in each tree considering a limited number of the variables. The final predictions of the random forest are made by averaging the predictions of each individual tree, which enhances the prediction accuracy for unseen data. The number of regression trees chosen for present modeling is (ntree) 500.

### Support Vector Machine

In Support Vector Machine, each data point is plotted in an n-dimensional space with the value of each variable being the value of particular coordinates and classification is performed based on the hyperplane that differentiates the two data classes. Following this, characteristics of new instances can be used to predict the class to which a new instance should belong.

### k-Nearest Neighbor

k-Nearest Neighbor (kNN) is one of the most basic and non-parametric algorithms, it does not make any assumptions about the distribution of the underlying data. The algorithm is based on the principle of Euclidean distance that is the instances within a dataset generally exist in close proximity to other instances that have similar properties. If the instances are tagged with a classification label, then the value of the label of an unclassified instance can be determined by observing the class of its nearest neighbors. For the present modeling, the whole process is repeated three times (repeats=3) each with a k value of 10 (number=10) and taking the average of the three iterations.

### Artificial Neural Network

An artificial neuron network (ANN) is based on the structure and functions of biological neural networks—the network learns (or changes) based on the input and output. The units in ANN are segregated into three classes: input units, which receive information to be processed; output units, where the results of the processing are found; and units in between known as hidden units. The network is first trained on a dataset of paired data to determine input-output mapping. The weights of the connections between neurons are then fixed and the network is used to determine the classifications of a new set of data. For the present modeling we consider the hidden units to be three (hidden=3) and threshold to be 0.05 (threshold=0.05).

To overcome our inability of using real-world data, we split the dataset into a “training set” (70%, i.e., 208 observations) and a “test set” (30%, i.e., 87 observations) making sure to balance the class distributions within the split (see the associated computer code on GitHub, https://github.com/aa54/CAD_1). The “training” dataset is used to train the model; the model sees and learns from this training data. The “test” dataset is then used to provide an unbiased evaluation of a final model fit to the training dataset. In some cases, we ran multiple experiments to validate model results on different splitting ratios.

The following four metrics[17] were used to evaluate the performance of the predicted models and compare them amongst one another.

1. Accuracy: the proportion of total dataset instances that were correctly predicted out of the total instances

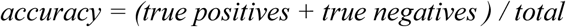
2. Recall (sensitivity): the proportion of the predicted positive dataset instances out of the actual positive instances;

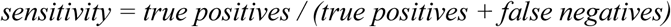
3. F1 score: a composite harmonic mean (average of reciprocals) that combines both precision and recall. For this, we first measure the precision, the ability of the model to identify only the relevant dataset instances

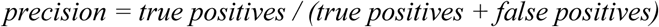 The F1 score is estimated as

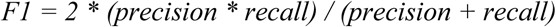
4. AUC (area under the curve): an estimate of the probability that a model ranks a randomly chosen positive instance higher than a randomly chosen negative instance. Additionally, the full AUC-ROC (Area Under the Curve–Receiver Operating Characteristic) curves are plotted to visually compare the models’ performance.

## Results and Discussion

For our machine learning model building, we started with two basic models, Logistic Regression and Decision Tree, to predict the presence of CAD. Our intuition was that it would be easy to interpret the results of the basic models and to explain model to the non-machine learning audience. After analyzing the results, however, we realized that the Decision Tree model is prone to over-fitting. and Logistic Regression did not perform relatively well with this simple dataset. We then considered applying the standard SMOTE methodology[18] to create additional synthetic data. Although we achieved more accurate results with SMOTE (5% to 6% higher accuracy), we felt that SMOTE is not necessarily a good approach as it creates data points based on the distance algorithm, and these synthetic datapoints may not be “true” representations of patients. Therefore, we decided not to perform SMOTE on the dataset and proceeded to apply the remaining, more complex ML algorithms.

As shown in Table 2, an accuracy value higher than 0.84 is achieved with all but the Decision Tree model (just below 0.80). The other performance parameters, recall, F1 score, and AUC are also high. A mean value is calculated with all the later three parameters for each of the models (shown in the last column in Table 2) to judge which model performs best as a whole. We excluded accuracy in calculating the Mean, as accuracy is often considered to be a misleading indicator in measuring the performance of models with biomedical datasets (e.g., see [3]).

**Table 2:** Model performance metrics

| Model | Accuracy | Recall | F1 score | AUC | Mean |
| --- | --- | --- | --- | --- | --- |
| Generalized Linear Model | 0.8764 | 0.8000 | 0.8786 | 0.883 | 0.85 |
| Decision Tree | 0.7978 | 0.7447 | 0.7970 | 0.801 | 0.78 |
| Random Forest | 0.8764 | 0.8261 | 0.8751 | 0.880 | 0.86 |
| Support-Vector Machine | 0.8652 | 0.7959 | 0.8662 | 0.871 | 0.84 |
| **Neural Network** | **0.9303** | **0.9380** | **0.8984** | **0.796** | **0.88** |
| k-Nearest Neighbors | 0.8427 | 0.7872 | 0.8419 | 0.847 | 0.83 |

As seen in Table 2, the performance of the Neural Network model is outstanding with an accuracy of 0.9303 and a recall of 0.9380. In addition, running multiple experiments with differently proportioned training and test sets (changing from 70:30 to either 60:40 or 80:20) with Neural Network, the performance metrics were unchanged.

A high recall value indicates a lower propensity for false negatives. In disease prediction, a low recall (high frequency of false negatives), would misdiagnose patients with CAD as healthy, which may have devastating consequences. For example, a recall rate of 0.9380 (achieved with Neural Network) implies that this algorithm will correctly identify the presence of CAD in approximately 94 patients out of 100 patients with CAD.

Figure 2 compares the performance of the six ML algorithms using AUC-ROC. AUC-ROC curves allow to visualize the tradeoff between the True Positive Rate and False Positive Rate, whereas AUC (see above) is useful to compare multiple algorithms or hyperparameters combinations (these two are obtained by different methods). As seen in the figure, all models except for Decision Tree perform well, there is very little difference in the AUC-ROC cures. Nevertheless, the ROC curve corresponding to the Neural Network model has a relatively higher positive slope, indicating that this model gives higher “true positive percentage.” Combining with the results obtained for the performance metrics, it may be concluded that the Neural Network model is able to predict CAD more effectively than the other models.

**Figure 2:**
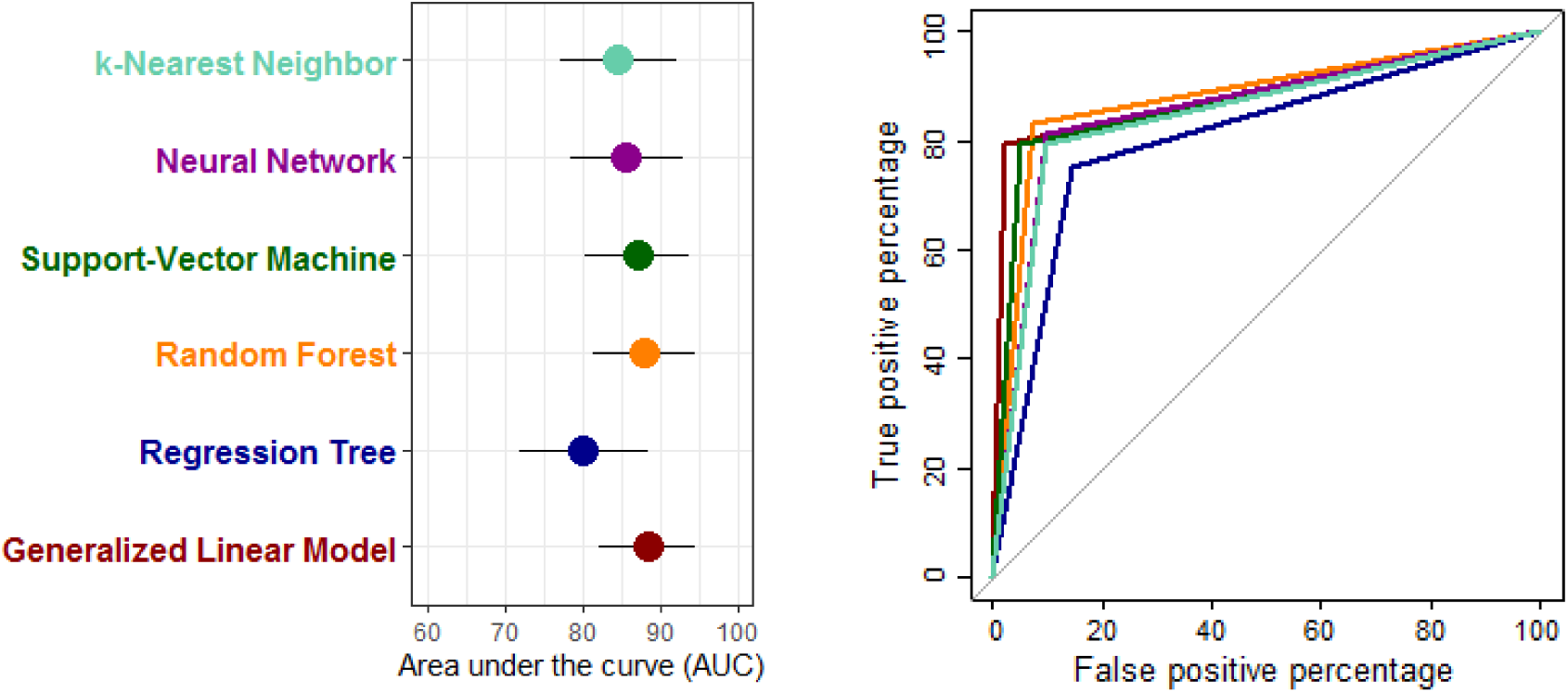
Classifier performance comparison with Receiver Operating Characteristic (ROC) curves. *Left:* Area under the ROC curve (AUC) for all the six models with estimated 95% confidence intervals. *Right*: Full ROC curves for the six models. Note the matching color code.

We next extracted at the “variable of importance” in the Neural Network model. Understanding the relative importance of variables may dictate which variables are necessary to be included in the risk prediction of CAD. As shown in Figure 3, “restcg” (resting EKG), and sex (gender) have a relatively high importance, while the three attributes “age,” “thalch” (maximum heart rate achieved during the thallium test), and “exang” (exercise induced angina) have the least importance. The top ten variables of importance as assigned by the Neural Network model, in descending order, are:

> resting EKG,
>
> patient gender,
>
> chest pain,
>
> slope of the peak exercise ST segment,
>
> ST depression induced by exercise relative to rest,
>
> fasting blood sugar (higher or lower than 120 mg/dl),
>
> thallium heart scan,
>
> number of major vessels,
>
> resting blood pressure, and
>
> serum cholesterol

**Figure 3:**
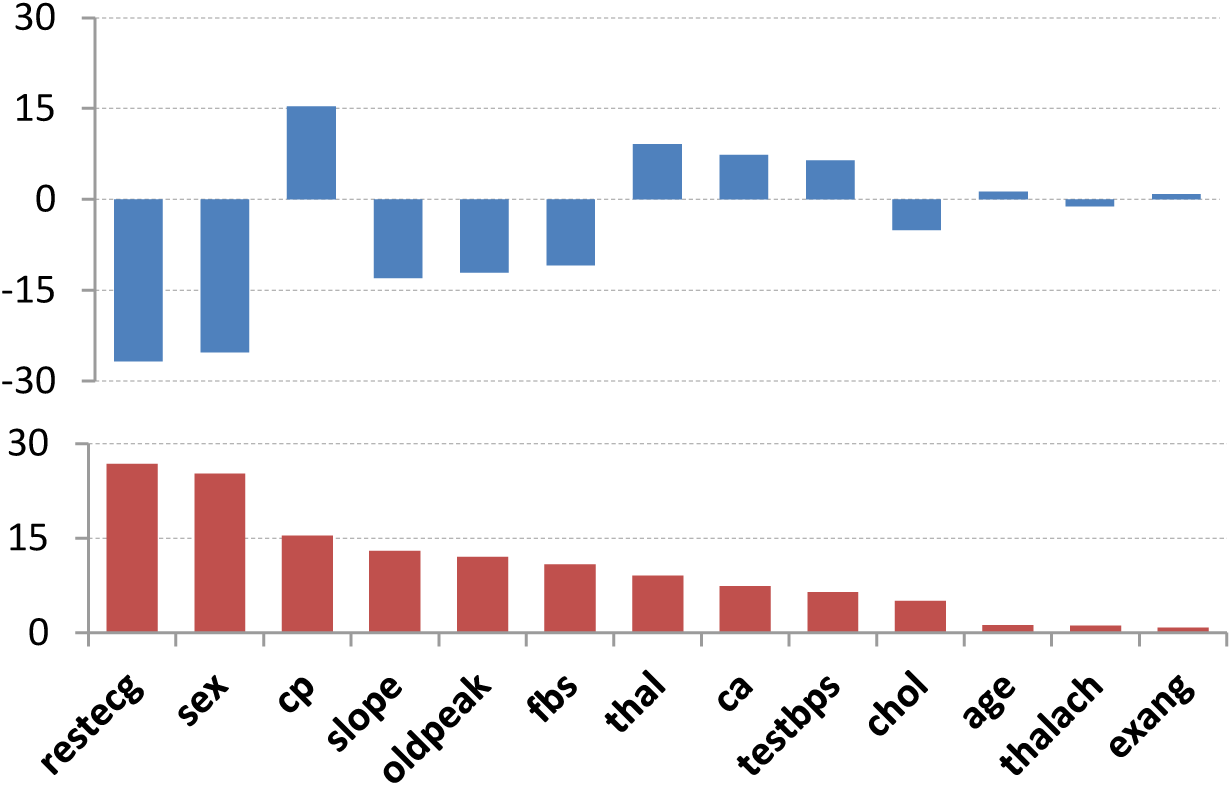
Variable of importance in the Neural Networks model. The *top* graph shows the importance as obtained with a built-in code library functiuon and the *bottom* is the normalized graph.

Heinze et al[19] proposed certain recommendations on the application of variable selection methods to help modeling in life sciences and its worth following these recommendations in future efforts of identifying which of the variables of importance are worth studying in the risk modeling.

## Conclusions and Future Scope

In this study, we demonstrated that ML algorithms can be applied with high accuracy and recall to detect the presence of CAD using a publicly available dataset. We also demonstrated that the Neural Network model outperforms other ML models to detect CAD. We deposited the associated computer code in the public domain (see https://github.com/aa54/CAD_1) in the hopes that we are contributing to an open source community.

**Figure 2:**
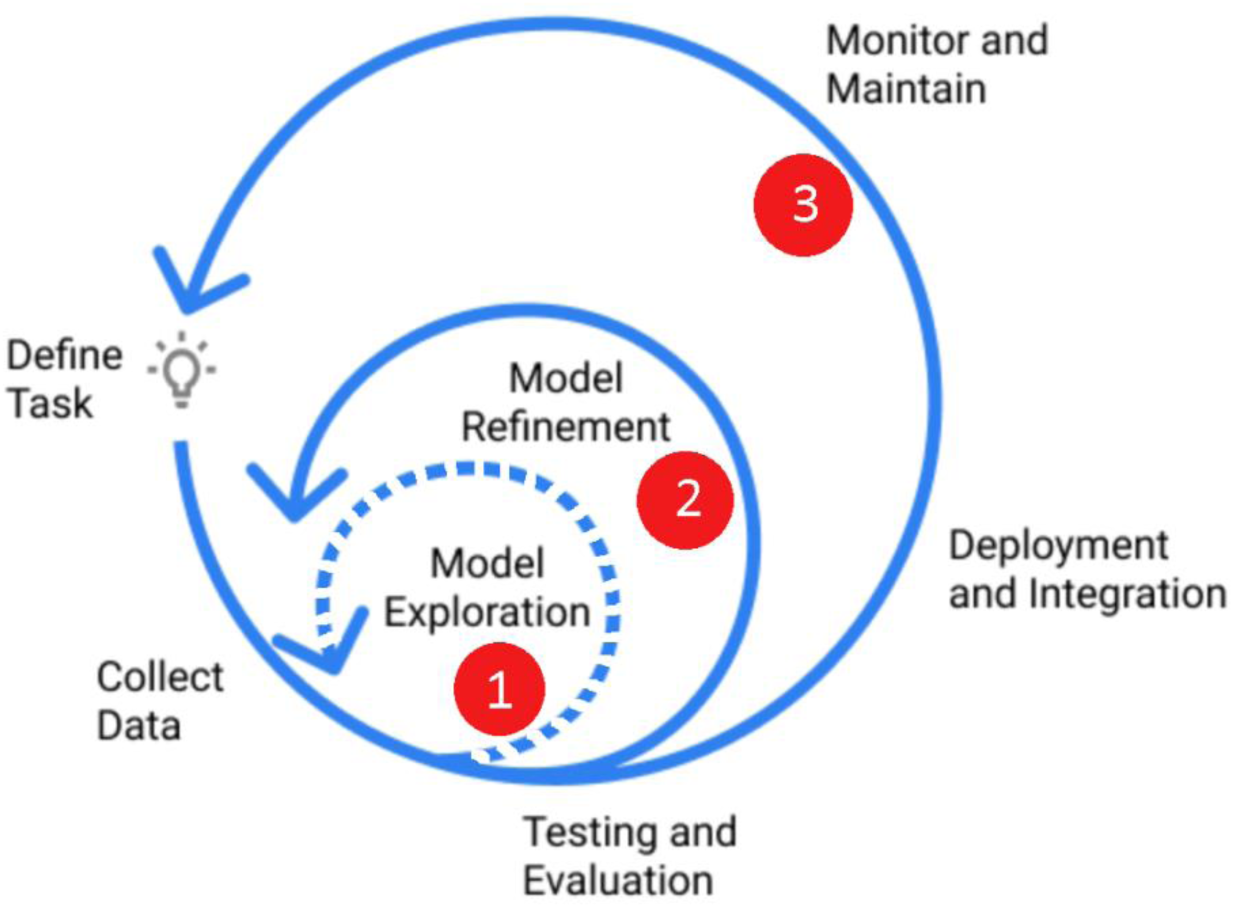
An ideal framework for developing a viable machine learning model. The dashed arrow indicates the iteration involving the ML algorithms presented here (also marked 1). Steps 2 and 3 (solid arrows) indicate future explorations. (Modified from https://www.jeremyjordan.me/ml-projects-guide/).

We visualize that a viable ML solution for predicting coronary artery disease (CAD) evolves in three steps: (1) model exploration; (2) model refinement; and (3) monitoring and maintenance. A proposed framework for arriving at a practical ML solution for the detection of CAD is depicted in Figure 4. We believe that with our current effort, we have achieved the first step (marked 1 in Fig. 4). We are hopeful that this study can help form the basis for further testing/validating of our algorithms with multiple and larger datasets. We look forward to gain access to larger datasets for further validation and refinement (depicted as step 2 in Fig. 4), with the eventual goal of providing an open source solution (step 3 in Fig. 4) to aid health care practitioners in the detection and treatment of CAD.

## Supporting information

Supplemental Material Data Analysis

## Footnotes

1 We noticed that a second copy of the “Cleveland dataset” is available on the UCI repository with a different name as “Statlog (Heart) Dataset” (see http://archive.ics.uci.edu/ml/datasets/statlog+(heart)) and several publications combine these two datasets for model building, or build on one dataset and validate on the other, which may lead to significantly misleading predictions.

