## Supplemental Material Data Analysis for "Machine Learning Algorithms for Predicting Coronary Artery Disease: Efforts Toward an Open Source Solution"

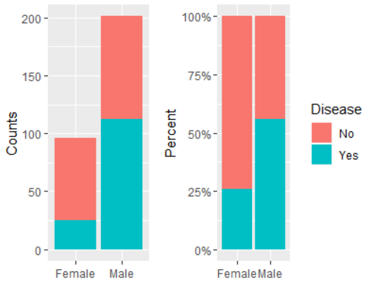


**S1:** Patients with and without heart disease.

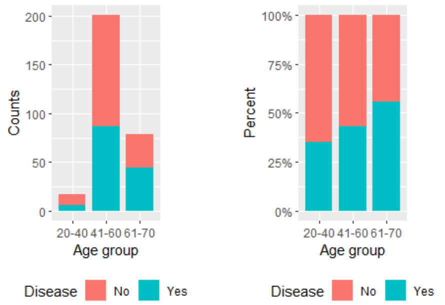


**S2:** Age of patients with and without heart disease.

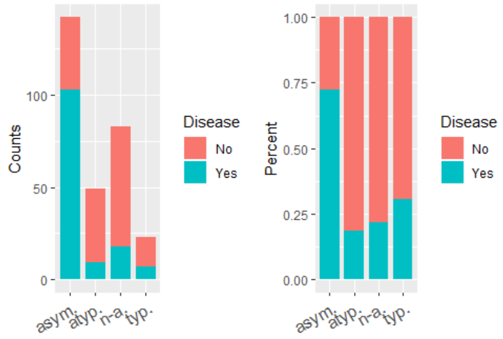


**S3:** Type of chest pain in patients with and without heart disease. Abbreviations: typ., typical angina; atyp., atypical angina; n-a., non-anginal pain; asym., asymptotic

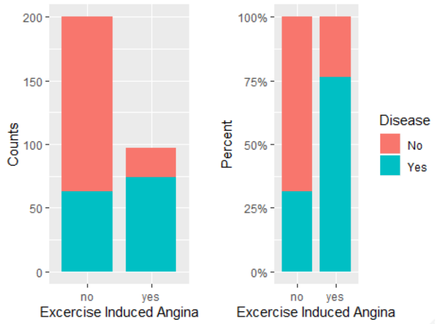


**S4:** Exercise induced angina (EIA) in patients with and without heart disease.

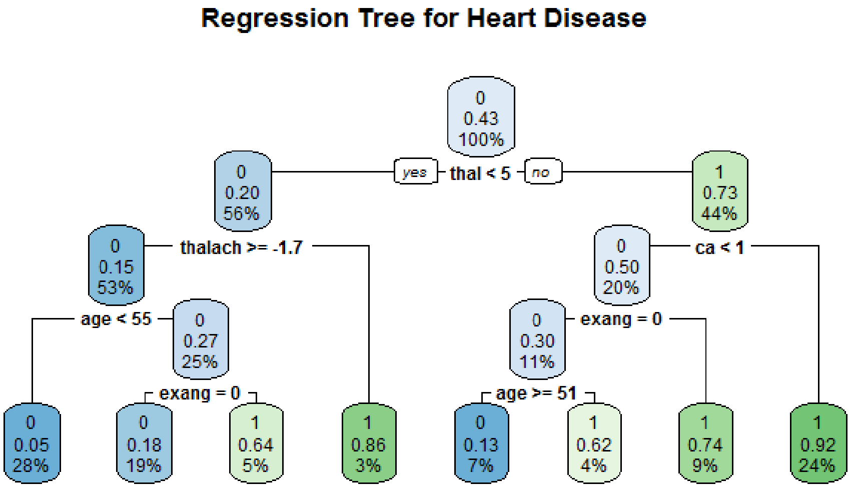


**S5:** Regression Tree Model

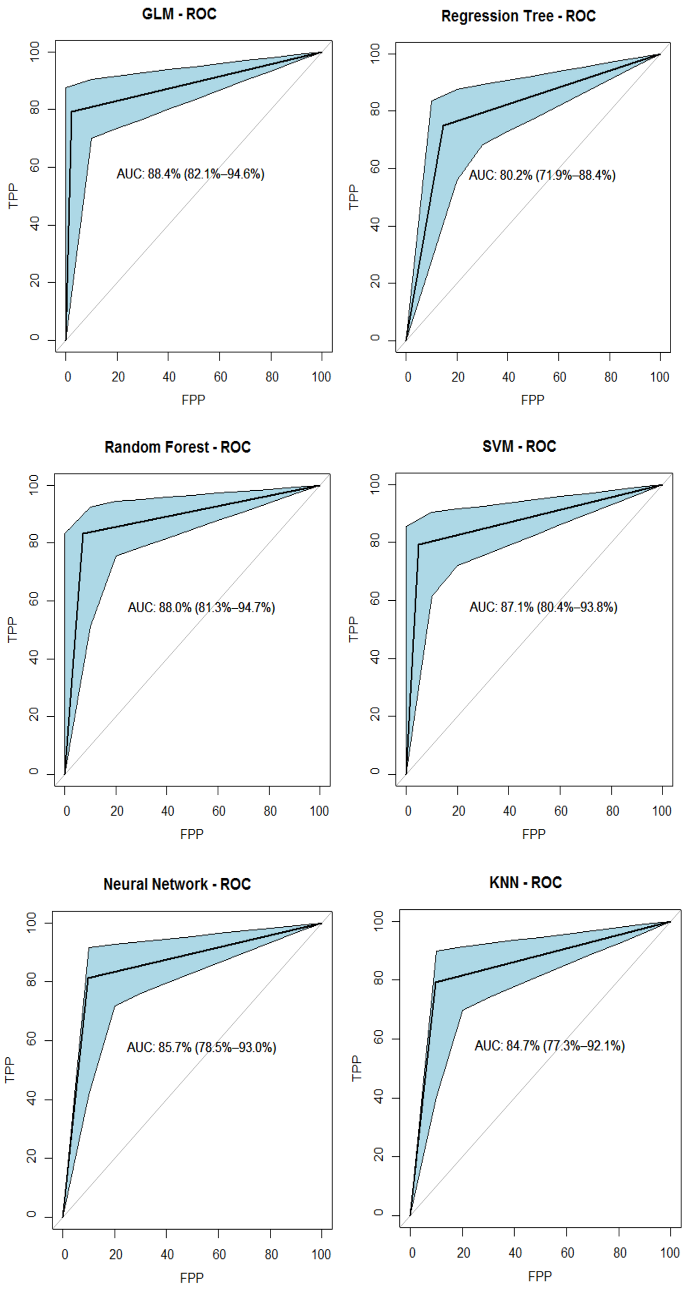


**S6:** Composite AUC-ROC curves
